# Language-aligned model and structured scene descriptions reveal sensitivity to compositional scene in the high-level visual cortex

**DOI:** 10.64898/2026.07.27.740322

**Authors:** Karim Rajaei, Arian Afshar, Radoslaw Martin Cichy, Hamid Soltanian-Zadeh

**Affiliations:** School of Cognitive Science, Institute for Research in Fundamental Sciences (IPM), Tehran 19568, Iran; Department of Biomedical Engineering, Amirkabir University of Technology, Tehran, Iran; Department of Education and Psychology, Freie Universität Berlin, Berlin, Germany; Control and Intelligence Processing Center of Excellence (CIPCE), School of Electrical and Computer Engineering, College of Engineering, University of Tehran, Tehran 14399, Iran

## Abstract

Natural scenes are defined not only by their contained objects, but also by the structured relations among those objects. Using 7T fMRI data from the Natural Scenes Dataset, we examined which information in scene-description representation contributes to the prediction of high-level visual cortical responses. We compared intact narratives with shuffled full description (bag-of-words), extracted content-word lists, and descriptions regenerated from those lists without access to the images of original narratives. Intact narrative descriptions outperformed the tested lexical controls. Both original and regenerated descriptions provided incremental predictive gains beyond bag-of-words, but the regenerated gain was only 21.3% as large as the original gain. Thus, constrained sentence reconstruction provided some additional predictive information without reproducing the original-description advantage. Exploratory image ratings indicated that this prediction advantage was greatest for scenes with richer scene compositional organization. Further, a language-aligned vision model provided additional predictive information beyond a self-supervised vision-only model, with a cortical distribution resembling the narrative advantage. These results address the representational basis of caption-based neural predictivity and are consistent with a contribution of structured scene information beyond simple entity co-occurrence. They further imply that language supervision may help induce comparable structure-sensitive representations in artificial vision models.

## Introduction

Natural scenes are not just collections of objects; they contain structured information about how entities are organized and interact, including spatial layout (Bar, 2004; Oliva & Torralba, 2007), semantic associations (Peelen et al., 2024), social interactions (Malik & Işık, 2023), and actions (Hafri et al., 2017). For example, the presence of “person,” “dog,” and “chasing” is not equivalent to representing whether the person is chasing the dog or the dog is chasing the person. Distinguishing these interpretations requires extracting relational structure, not just detecting individual entities. The visual system rapidly extracts such structural information, allowing coherent interpretation of scene meaning beyond local visual features (Hafri & Firestone, 2021; Peelen et al., 2024; Wischnewski & Peelen, 2021). Such scene understanding also depends on prior knowledge that constrains and organizes perceptual interpretation (Aronson et al., 2025; Collins & Olson, 2014; Võ, 2021).

Here, we focus on compositional structure: the meaningful organization of scene elements— agents, objects, actions, and spatial layout—into a coherent whole (Hafri & Firestone, 2021; Lake et al., 2017; Malik & Işık, 2023). Natural scenes are therefore defined not only by which elements are present, but also by the relations that link them, including spatial, action-based, and social relations; entity co-occurrence describes which elements appear together without explicitly specifying these relations (Greene, 2013). We hypothesize that representation preserving the organization of scene descriptions may provide information relevant to high-level visual responses beyond the representations based on lexical content alone. Natural language provides a useful probe of such structure because it explicitly encodes relations among objects, actions, and spatial configurations in a structured form (Cavanagh, 2021; Jackendoff, 1985).

Previous studies have established that caption-derived representations predict high-level visual responses, but they also raised a more specific question: which information within those representations drives neural predictivity? Doerig et al. (2025) found that intact and scrambled captions produced highly similar embeddings. They suggested that the encoder might recover meaning from simple scrambled NSD captions and explicitly left open whether this would hold for more complex descriptions in which word order matters. Conwell et al. (2024) reported that predictivity was driven predominantly by nouns, with little contribution from syntax or semantic compositionality. These findings motivate testing the predictive contribution associated with description organization beyond the lexical controls used here. Our study addresses this question using detailed generated descriptions, shuffled full descriptions, extracted content-word lists, and constrained regeneration within a common encoding framework. The contribution is to characterize the information supporting caption-based predictivity, rather than to establish caption–brain alignment itself. We also examined whether selected images associated with the model differences have richer relational or compositional structure.

The control address successive questions. First, comparing original descriptions with bag-of-words tests whether their original organization provides predictive information beyond the same input word inventory in randomized order. However, scrambling also disrupts fluency and grammatical or sequential form, so an original-over-bag-of-words advantage alone does not identify image-related compositional information. Second, we regenerate descriptions from extracted content-word lists without providing the image or original narrative. This procedure is intended to restore grammatical sentence form while reducing recovery of the original image-specific organization. Comparing regenerated descriptions with bag-of-words asks whether newly constructed sentence-like text provides additional predictive information beyond shuffled full descriptions. Comparing original with regenerated descriptions ask whether the original description retain an advantage over that reconstruction procedure.

Second, we complement the description-based analyses with a model-level comparison between SigLIP2 and DINOv3, two large vision-transformer models trained with language-aligned and self-supervised visual learning objectives, respectively (Chen et al., 2020; Radford et al., 2021). Previous studies have reported that language-aligned representations can improve prediction of visual responses (Doerig et al., 2024; Rajaei et al., 2026; Wang et al., 2023), but the basis of this advantage remains unclear. This comparison asks whether language alignment is associated with neural predictivity that resembles the information retained by intact scene descriptions. Specifically, we test whether SigLIP2 provides additional predictive information relative to DINOv3, and whether the spatial distribution of this difference resembles the advantage associated with intact narrative descriptions.

## Results

### Narrative scene descriptions outperform lexical controls in predicting cortical responses

We first asked whether structured narrative descriptions outperform lexical control representations in predicting cortical responses to natural scenes. Using high-resolution 7T fMRI data from the Natural Scenes Dataset (NSD), we fit voxelwise encoding models to data from eight participants, each of whom viewed 10,000 unique natural images. For each image, we generated a narrative description using a generative vision-language model (VLM). From these descriptions, we derived multiple linguistic feature spaces that varied in syntactic and lexical structure, including full narrative descriptions, bag-of-words representations (same description words in shuffled order), content words (extracted lists of nouns, verbs, and adjectives), and isolated lexical categories of nouns, verbs, and adjectives. An additional regenerated-description control reconstructed descriptions from content-word lists without access to the images. These representations were embedded with a text encoder and used to predict held-out fMRI responses in cross-validated voxelwise models (Fig. 1a). Unless otherwise noted, results are reported for embeddings derived from the CLIP text encoder; analyses using a language-only RoBERTa encoder yielded a closely similar pattern of results (Fig. 1c), indicating that the effects were not specific to a particular text-embedding architecture.

**Figure 1.**
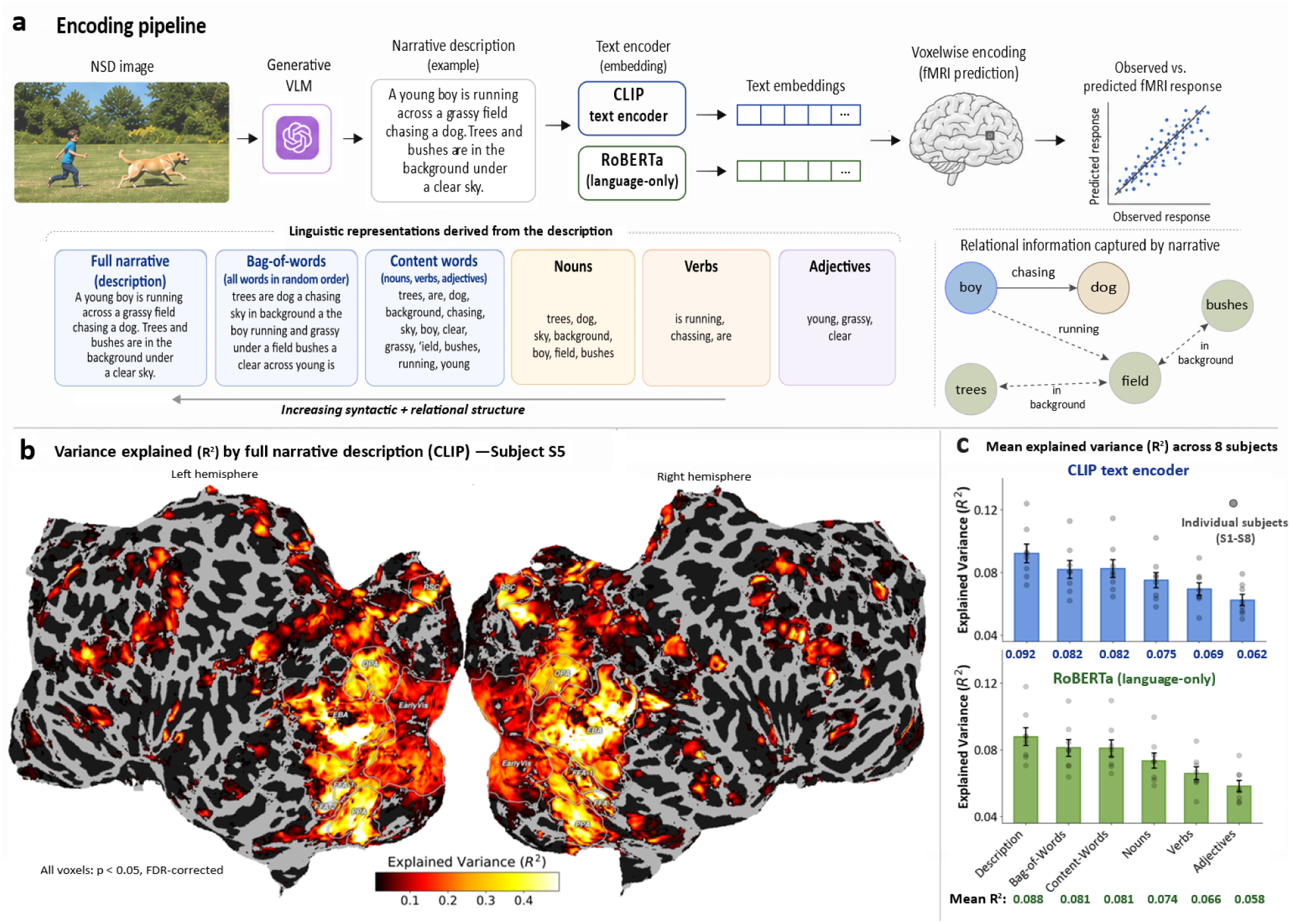
Variance explained by language-based scene descriptions. **a.** Schematic of the encoding pipeline. For each NSD image, a VLM produces a narrative description. Text embeddings derived from these descriptions are then used in voxelwise encoding models to predict fMRI responses. **b.** Variance explained (R²) by the narrative description on a cortical flatmap for a representative subject S5 (all subjects in Supplementary Fig. 1). Only significant voxels are shown (bootstrap test, p < 0.05, FDR-corrected). **c.** Mean explained variance (R²) across subjects for full narrative descriptions, bag-of-words, content words, and isolated lexical categories (objects, verbs, and adjectives). Bag-of-words denotes shuffled full descriptions, content words denotes extracted word lists. Error bars denote standard deviation; dots indicate individual subjects.

Full narrative descriptions significantly predicted activity across broad portions of visual cortex (Fig. 1b). Prediction performance was evident throughout occipital and ventral temporal cortex and was strongest in higher-level category-selective regions, including EBA, FFA, PPA, OPA, and RSC. Performance was lower in the early visual cortex, consistent with the view that narrative embeddings align more strongly with higher-level scene representation than low-level visual features (Doerig et al., 2024; Rong et al., 2025; Wang et al., 2023).

To isolate the predictive contribution associated with description organization, we compared full narratives with bag-of-words. This contrast retains the original words while randomizing their order. Across subjects, full narratives significantly outperformed bag-of-words (p < 0.01, Wilcoxon signed-rank test; Fig. 1c), demonstrating that intact descriptions provided predictive information beyond the presence of scene-related words alone. Equal word inventories at the input do not establish equal lexical information in the embeddings, so this difference cannot be attributed uniquely to compositional structure.

To examine the effect of retaining function words in unstructured inputs, we compared bag-of-words with content-words lists. Bag-of-words retains the words of the full description, including function words, such as prepositions, determiners, and conjunctions, whereas the content-words procedure extracted a reduced word list. We did not detect a significant difference in prediction performance between these conditions (Fig. 1c; p > 0.05). This result does not establish equivalence between them or rule out a contribution of function words.

We next examined which lexical categories carried the most information by comparing models based on nouns, verbs, or adjectives alone. Nouns explained the most variance, followed by verbs and then adjectives, with all pairwise differences significant (p < 0.05; FDR-corrected). However, this ordering was partly attributable to differences in word counts across categories. After matching word counts across nouns, verbs, and adjectives to the minimum available count, the difference between nouns and verbs was no longer significant, whereas adjectives still explained significantly less than both nouns and verbs (Supplementary Fig. 2). All single-category models performed worse than the content-words baseline (p < 0.01; FDR-corrected), indicating that unordered combinations of multiple lexical categories carry more predictive information than any category in isolation. Because this comparison varies both lexical coverage and input length, however, it does not provide a clean test of whether the gain arises specifically from combining categories.Together, these results show an advantage for intact descriptions over tested lexical controls. We next examined incremental predictivity and the regenerated description control to characterize this advantage further.

### Language-aligned and narrative representations show similar distribution of incremental predictivity

We next asked where intact descriptions explain variance beyond bag-of-words representations. Using joint voxelwise encoding models that included both narrative and bag-of-words features, we computed incremental explained variance, termed unique R² here (Fig. 2a). This contrast quantifies the gain from adding narrative features to the tested baseline within the specified ridge-regression pipeline. Narrative descriptions showed significant unique R² concentrated in high-level category-selective regions of ventral temporal and lateral occipital cortex, with little or no unique variance in early visual cortex. The reverse contrast (bag-of-words > narrative) yielded no significant voxels. Group-level ROI analyses showed significant unique R² in body-, face-, and place-selective regions (*all p* < 0.01), with smaller but nonetheless significant unique R² in early visual cortex (Fig. 2b).

**Figure 2.**
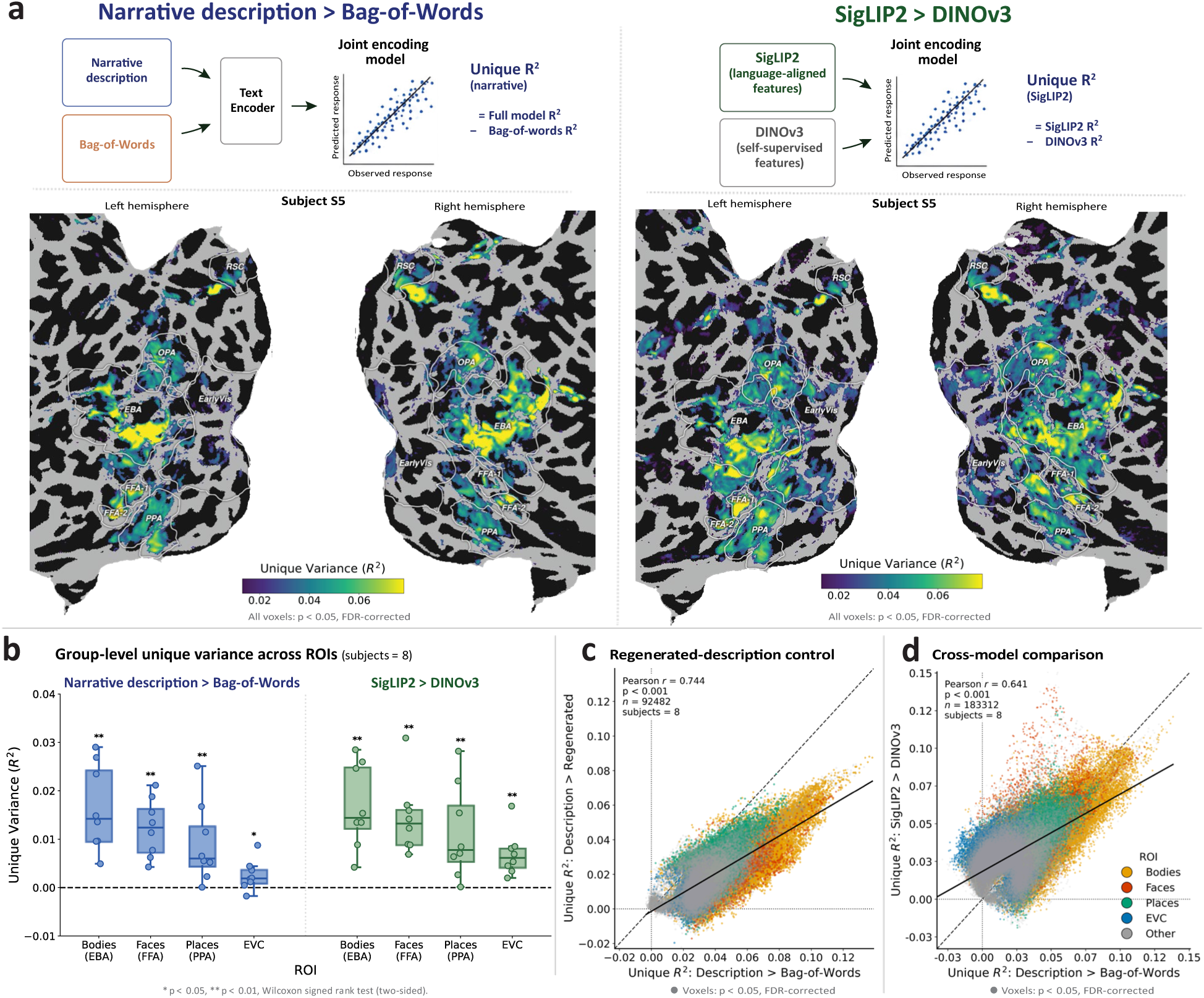
Unique variance explained by narrative descriptions, regenerated-description control, and language-aligned visual model. **a.** Voxel-wise unique explained variance (R²) for narrative descriptions relative to bag-of-words (left) and for SigLIP2 relative to DINOv3 (right), shown for subject S5 as an illustrative visualization (all subjects in Supplementary Fig. 3). In each joint encoding model, both feature sets are entered simultaneously; unique R² is defined operationally as the joint-mode R² minus baseline-model R². Only significant voxels are displayed. **b.** Group-level unique R² across ROIs for narrative description > bag-of-words (left) and for SigLIP2 > DINOv3 (right). Box plots show the median and interquartile range across eight subjects; points indicate individual subjects. Asterisks denote significance against zero (two-sided Wilcoxon signed-rank test, * p < 0.05, ** p < 0.01). **c.** Regenerated-description control in eight subjects. Each point represents a voxel. The x-axis shows unique variance for original narrative descriptions relative to bag-of-words, and the y-axis shows unique variance for original narrative descriptions relative to regenerated descriptions. Text inset reports the voxelwise correlation, p-value, and number of voxels, pooled across subjects. Only voxels with significant unique variance in either contrast are shown (p < 0.05, FDR-corrected). **d.** Correlation between unique variance explained by description and vision models across cortical ROIs. Each point represents a voxel showing significant effects in either contrast, pooled across subjects. The x-axis shows unique variance for description > bag-of-words, and the y-axis shows unique variance for SigLIP2 > DINOv3. Colors denote ROIs: body-selective cortex, face-selective cortex, place-selective cortex, and early visual cortex. Across included voxels, the two contrasts are positively correlated (r = 0.64, p < 0.001), and the solid line indicates the best-fitting linear regression.

To distinguish the original narrative advantage from a benefit of having sentence-like input, we performed a regenerated-description control in all eight subjects. The bag-of-words condition preserves the full original word inventory but disrupts its order and grammatical form. We therefore supplied extracted content-word lists to a language model without the image or original narrative and instructed it to construct a grammatical description using those words and only necessary function words (Supplementary Table 1).

We quantified the incremental gain of each description representation beyond the same bag-of-words baseline. Original descriptions provided a gain of 0.03969 R² units (95% CI [0.03746, 0.04164]). Regenerated descriptions also provided a positive gain (0.00846 R² units, 95% CI [0.00590, 0.01092]; two-sided subject-level sign-flip test, p = 0.008), but the original-description gain was larger (paired difference = 0.03124 R² units, 95% CI [0.02822, 0.03444]; two-sided paired subject-level sign-flip test, p = 0.01). The regenerated gain was 21.3% as large as the original gain (95% CI [14.8%, 27.4%]). Thus, constrained regeneration provided additional predictive information beyond bag-of-words but did not reproduce the magnitude of the original-description advantage. The ratio describes relative gain magnitude, not the fraction of shared variance or a decomposition into grammatical and image-specific information.

Separately, we compared original and regenerated descriptions using the same variance-partitioning procedure. Across voxels, the incremental R² for original description beyond regenerated descriptions was correlated with the incremental R² for original description beyond bag-of-words (r = 0.744, p < 0.001; Fig. 2c; see Supplementary Fig. 4 for S5). Within the common voxel set, the median original-versus-regenerated incremental R² was 66% of the median original-over-bag-of-words incremental R². This is a different quantity from the regenerated-over-bag-of-words gain ratio reported above, it compares original-description gains conditional on two different baselines.

We quantified this residual similarity across 73,000 aligned original–regenerated description pairs. As intended, the pairs had nearly identical non-function-word inventories (mean token-overlap F1 = 0.957). However, longer verbatim sequences were rare: the median longest shared sequence was two non-function words, and only 11.3% and 1.6% of pairs shared sequences of at least three and four words, respectively. Word-order similarity was also only modestly greater than a within-pair null in which the same regenerated content words were randomly reordered (mean ROUGE-L: 0.446 vs. 0.405; difference = 0.041, 95% CI [0.040, 0.041]; Supplementary Table 2). These measures document substantial lexical preservation with limited extended verbatim copying. They do not establish matched fluency or that the regenerated descriptions express relations different from those in the image. The original-over-regenerated encoding comparison nevertheless demonstrates additional predictive information beyond the representations produced by this reconstruction procedure.

We then compared SigLIP2, a language-aligned model, with DINOv3, a self-supervised vision-only model from the same broad architecture family, both trained at internet scale. Unique R² for SigLIP2 relative to DINOv3 showed a distribution similar to the narrative-over-bag-of-words result, localizing to high-level category-selective cortex with minimal effects in early visual areas (Fig. 2a). The reverse contrast yielded no significant voxels. ROI analyses showed significant unique R² across category-selective regions (all p < 0.05; Fig. 2b).

We next asked whether the same voxels showed incremental predictivity for intact descriptions and language-aligned visual features. Across significant voxels pooled across subjects (n = 183,312), unique variance for narrative descriptions over bag-of-words was correlated with unique variance for SigLIP2 over DINOv3 (r = 0.641, p < 0.001; Fig. 2d). Positive correlations were observed in body-selective (r = 0.68), face-selective (r = 0.60), and place-selective cortex (r = 0.54), where the distribution was shifted toward greater unique variance for narrative descriptions than for SigLIP2. In early visual cortex, however, the correlation was weakly negative (r = −0.10), and the distribution was shifted toward greater unique variance for SigLIP2 (all p < 0.001; see Supplementary Table 3 for subject-level statistics). These results describe spatial correspondence between the two contrasts within the selected voxel set. They do not establish identical representational content.

### Exploratory characterization of images with high prediction-difference alignment scores

To characterize images associated with the model differences, we computed a prediction-difference alignment score across significant voxels for each contrast (see Methods). This score uses observed responses and joint-minus-baseline predictions, it is not an exact per-image decompositions of unique R2. For each contrast, we selected the 20 images with the largest positive scores and 20 images with scores closest to zero. These images were used for the quantitative analyses and box plots; 10 high scored images from each group were displayed for visualization.

We then annotated each selected image using three descriptive measures: compositional structure, semantic relations, and social interaction. For the description > bag-of-words contrast, high-score images scored higher than near-zero images on all three descriptive measures: compositional structure (9.25 ± 2.24 vs. 5.20 ± 1.96; p < 0.001), semantic relations (5.45 ± 1.47 vs. 2.35 ± 1.09; p < 0.001), and social interaction (2.25 ± 0.85 vs. 0.35 ± 0.59; p < 0.001; Fig. 3). The same pattern was observed for SigLIP2 > DINOv3. Highscore images again had higher compositional structure scores (8.55 ± 3.27 vs. 6.20 ± 2.28; p = 0.007), semantic relational scores (5.00 ± 1.86 vs. 3.15 ± 1.18; p = 0.0013), and social interaction scores (1.40 ± 1.10 vs. 0.30 ± 0.73; p = 0.001). Ratings for all selected images are shown in Supplementary Fig. 5.

**Fig. 3.**
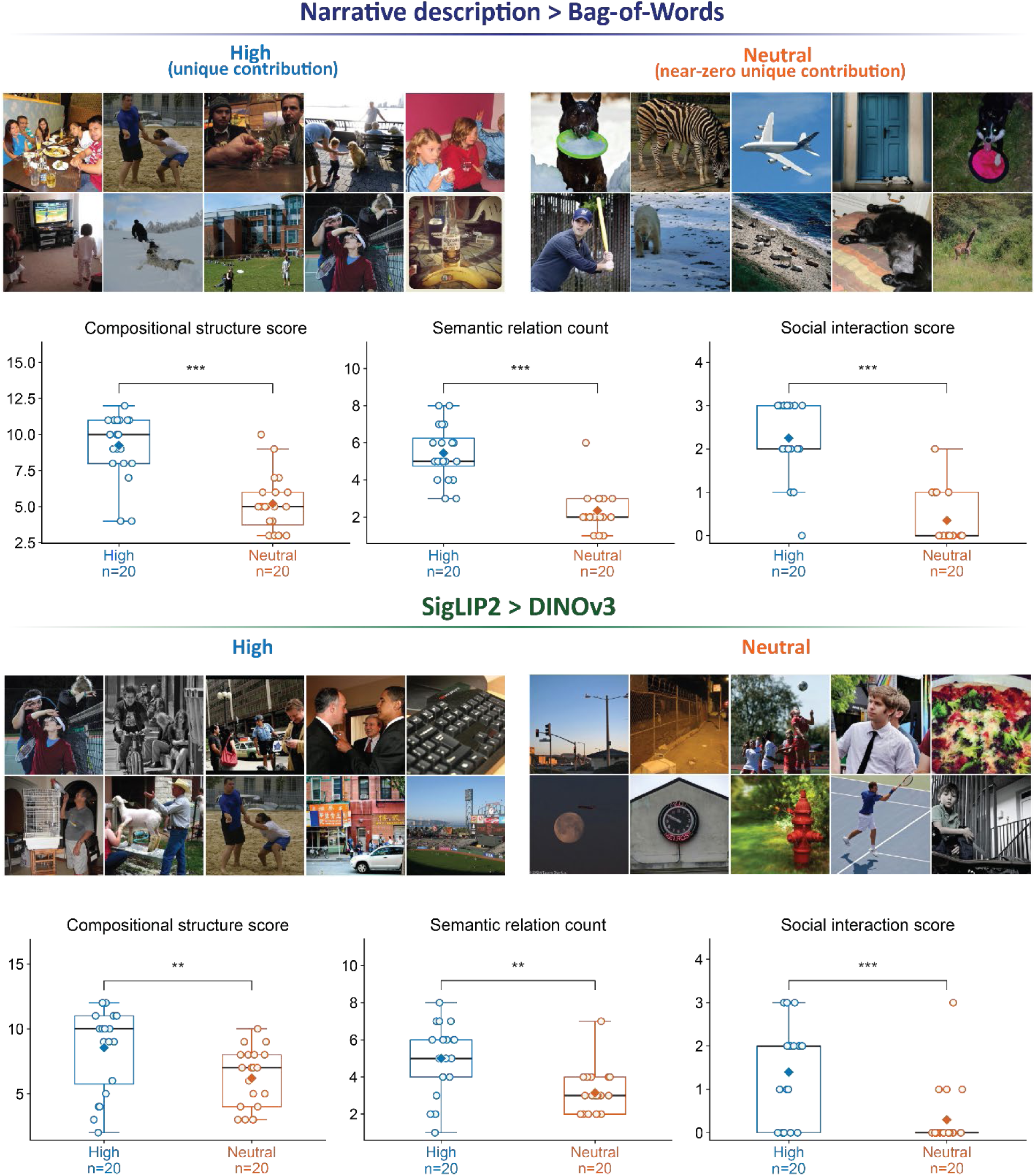
Stimulus-level drivers of unique variance. Top, narrative description > bag-of-words: the 10 displayed examples from the high-score group are shown on the left, and 10 examples from the near-zero group (labeled neutral) are shown on the right. Bottom, SigLIP2 > DINOv3: corresponding images groups for the visual model contrast. The image-ranking score is defined in Method and is distinct from voxelwise unique R^2^. Box plots summarize three descriptive compositional structure, semantic relational, and social interaction scores for 20 images per group. Blue boxes indicate high-score images and orange boxes indicate near-zero images; circles show individual image ratings. Boxes show the median and interquartile range. Asterisks indicate differences between groups using two-sided Mann–Whitney U tests (*p < 0.05, **p < 0.01, ***p < 0.001).

Under the study-specific annotation scheme, high-score images received higher ratings of scene organization, visible semantic relations, and social interaction than near-zero images. This exploratory association does not distinguish relational organization from correlated properties such as agent number, entity diversity, or scene richness.

## Discussion

Our results address which information within scene-description representation supports high-level visual neural predictivity. Intact narrative descriptions outperformed the tested lexical controls and provided additional predictive information beyond descriptions regenerated from content-words lists. Regenerated descriptions also provided a positive gain beyond bag-of-words, but that gain was substantially smaller than the original-description gain. Thus, sentence reconstruction recovered some predictive information without reproducing the magnitude of the original advantage. The narrative advantage was concentrated in body-, face-, and place-selective regions and was smaller in early visual cortex, and was most pronounced for images with richer compositional structure. These findings are consistent with a contribution of structured scene information, while leaving its specific representational basis unresolved.

The distinction from prior work lies in the question tested and the pattern of results. Doerig et al. (2025) reported highly similar MPNet embeddings for intact and scrambled captions and proposed that meaning could be recovered from simple scrambled descriptions. Our comparisons examine the unresolved role of description organization using detailed narratives and complementary controls. Conwell et al. (2024) attributed predictivity largely to nouns, whereas we observed advantages for intact descriptions over the tested lexical representations and incomplete recovery of that advantage through regeneration. This difference limits generalization of a predominantly noun-based account to the descriptions, encoders, and encoding procedure examined here; it does not refute the earlier results under their own conditions. Caption content and encoder choice can influence predictivity (Bavaresco et al., 2026), and differences in these factors or evaluation procedures may contribute to the discrepancy. Our study therefore extends the investigation of what drives caption-based predictivity, without claiming the first evidence of language–vision alignment or establishing sentence complexity as the explanation for cross-study differences.

The two controls support a sequential interpretation. The original-over-bag-of-words advantage establishes a benefit of the intact representation over shuffled input, but scrambling may impair lexical encoding, fluency, or sequential processing. Regeneration tests whether constructing new sentence-like text from the core content words can recover that benefit. Its positive incremental R² beyond bag-of-words is consistent with a contribution from restored sentence form, inferred semantic relations, or other changes introduced during generation. However, this gain was only 21.3% as large as the original-description gain. Generic reconstruction under the tested procedure was therefore insufficient to reproduce the original advantage. This percentage compares gain magnitudes; it does not partition the original gain into shared and image-specific components. In particular, the remaining 78.7% cannot be assigned uniquely to image-specific organization. The comparison of the two gains beyond bag-of-words also does not itself establish additional information conditional on regeneration; that question is addressed by the separate original-over-regenerated analysis.

Three candidate accounts help interpret this pattern. Under a lexical-content account, similar predictivity would be expected if regeneration preserved all relevant lexical information. Under a generic linguistic-form account, restoring adequate grammatical and sequential structure would be expected to recover much of the original advantage. An account emphasizing the original image-description organization predicts a remaining original advantage. The observed partial recovery and residual advantage are consistent with the latter account, but do not isolate it: the regenerated outputs also differ in fluency, wording, lexical realization, and generation style. They may infer image-consistent relations from the image-derived words even without access to the image. Low sequence overlap therefore does not demonstrate altered scene relations.

The comparison between SigLIP2 and DINOv3 provides a complementary result at the level of visual representations. Both models belong to the ViT-L family and were trained at internet scale, placing the comparison within a common broad architectural and large-data regime. SigLIP2 explained additional variance beyond DINOv3 in high-level category-selective cortex, and this advantage had a cortical distribution similar to the narrative advantage-over-bag-of-words. This convergence suggests that language-aligned training may emphasize representational dimensions that are also relevant to high-level visual cortex. Although this does not isolate the causal effect of language supervision.

The similar pattern observed across CLIP- and RoBERTa-based text embeddings indicates that the main narrative advantage was not restricted to one tested encoder. Instead, the results point to a broader correspondence between structured linguistic representations and high-level visual representations. In this context, language is not a substitute for vision. Rather, it provides a useful probe for isolating relational information that the visual cortex may itself encode.

Several considerations constrain interpretation. First, scrambling changes both sequential organization and familiarity of the input to the encoder; regeneration changes sentence form potentially its correspondence to the image. Neither manipulation was independently validated as a isolated change in compositional meaning. Second, the voxelwise correlations were computed after significance-based selection and pooling across participants. Spatial dependence, selection, and shared variation in prediction reliability may influence these associations. Finally, differences in raw predictivity across EBA, FFA, and PPA may reflect category selectivity, response reliability, or the information covered by descriptions; the present analyses do not distinguish these explanations or establish an ordering of relational sensitivity across regions.

An important next step is to decompose which kinds of relational information may contribute to cortical predictivity. Spatial configuration, action structure, physical interaction, agent-object binding, and social relations may make distinct contributions and may differ in their cortical distribution and functional relevance (Epstein & Baker, 2019; Hafri & Firestone, 2021; Isik et al., 2017; Kominsky & Wenig, 2025; Pitcher & Ungerleider, 2021; Wurm & Caramazza, 2019). Separating these components would help determine whether high-level visual cortex contains a common code for structured scene meaning or multiple specialized representations tuned to different relational dimensions.

A related question is whether scene-related representations are tied to visual input or generalize across modalities. For example, an image of a person chasing a dog, a sentence describing the same event, and a simple schematic depiction differ in sensory format but share a common relational structure. Future work could test whether high-level visual regions represent such shared relations across images, language, and schematic displays, building on evidence for partially modality-invariant semantic representations across visual and linguistic formats (Fairhall & Caramazza, 2013; Popham et al., 2021; Simanova et al., 2014; Wurm & Caramazza, 2019). Such studies would help determine whether predictive scene information observed here reflects visual scene-specific organization or a more abstract code for relations such as spatial layout, agent-action-patient structure, and social interaction.

More broadly, these results add to the growing use of generative vision-language models as tools for cognitive neuroscience (Luo et al., 2024; Panchal et al., 2026; Rong et al., 2025; Ye et al., 2025). Rich image-specific descriptions can be systematically perturbed while retaining overlapping word inventories. Such manipulation depends on the validity of the controls. This approach complements controlled stimulus manipulations by characterizing model predictivity for complex natural scenes.

Intact scene descriptions provided predictive information beyond shuffled-word and regenerated-description controls. Regeneration yielded a positive but substantially smaller gain beyond bag-of-words, showing that newly constructed sentence-like descriptions did not reproduce the full original-description advantage. Together with the complementary SigLIP2–DINOv3 comparison, these results refine the question of what information in language-related representations supports high-level visual neural predictivity. They are consistent with a role for structured scene information, but the contributions of compositional meaning, fluency, lexical encoding, and pretrained-model differences remain unresolved.

## Methods

### Dataset

We analyzed data from the publicly available Natural Scenes Dataset (NSD), a large-scale 7T fMRI dataset acquired from eight adult participants. Each participant completed up to 40 scanning sessions and viewed 10,000 unique natural images, yielding ∼73,000 image trials across subjects. Of the 10,000, 1,000 were shared across all participants to enable cross-subject comparisons. Stimuli were sampled from the Microsoft Common Objects in Context (COCO) dataset and consisted of complex natural scenes containing multiple objects embedded in structured contextual backgrounds.

Each image was presented for 3 s, followed by a 1 s inter-trial interval (4 s trial duration), subtending ∼8.4° of visual angle. Images were repeated up to three times within or across sessions. Participants performed a continuous recognition task, responding when an image was repeated.

Functional MRI data were acquired at 7T using gradient-echo echo-planner imaging (EPI) with 1.8 mm isotropic voxels and TR = 1.6 s. We used the publicly released single-trial beta estimates (betas_fithrf_GLMdenoise_RR), derived from a general linear model (GLM) that included (i) hemodynamic response function (HRF) fitting, (ii) GLMdenoise regressors to remove structured noise components, and (iii) ridge regularization to stabilize estimates. Beta weights were estimated independently for each voxel and trial.

For all encoding analyses, beta weights were z-scored voxel-wise within each session to reduce session-specific variance. For images presented multiple times, beta values were averaged across repetitions, yielding one response pattern per image per voxel.

All analyses were conducted in each subject’s native cortical surface space. Surface visualizations were generated using Pycortex.

### Regions of interest (ROIs)

ROI definitions were taken from the NSD release and included the early visual cortex and higher-level category-selective regions. Unless otherwise noted, left- and right-hemisphere ROI labels were merged within the subject prior to summary analyses.

### Stimulus annotation and linguistic feature spaces

To obtain detailed image-grounded descriptions, we generated scene annotations using Qwen 2.5-VL, a generative vision-language model. Inference was performed independently for each participant on 10,000 images.

We used a constrained prompt requesting a concise description (≤ 40 words). We distinguish six primary linguistic feature spaces and one regeneration control. Narrative description is original generated sentence description. Bag-of-words contains the words of that full description in shuffled order. Content-words is an extracted word list represent as [word1, word2, …], rather than a shuffled full description. Nouns (objects), verbs (actions), and adjectives (attributes) are the isolated lexical-category inputs. A regenerated description is a new description constructed from the content-word list without access to the image or original narrative.

The term bag-of-words thus names the input manipulation; after embedding with a transformer, the resulting representation is not guaranteed to be order-invariant. For the word-count control, noun, verb, and adjective inputs were matched to the minimum available count across categories, as reported in supplementary Fig. 2.

All textual inputs were embedded using the CLIP text encoder (ViT-L/14) and language-only RoBERTa. For each input, we extracted the model’s pooled text embedding and applied L2 normalization.

### Regenerated-description control

To complement the bag-of-words comparison, we performed a regenerated-description control in all eight subjects. The aim was to test whether restoring grammatical and sequential form to scene-related content words could recover the original narrative advantage. For each original generated description, we removed function words to obtain a content-word list containing the scene-relevant nouns, verbs, and adjectives. These words were provided in randomized order to GPT-5.5 through Codex, without the image or original narrative. Initial regeneration attempts using Qwen produced unsatisfactory outputs, motivating the use of GPT-5.5 for this separate text-only task. The model was instructed to generate one grammatical description using all and only the supplied content words, while adding only necessary function words, such as determiners, auxiliaries, conjunctions, and prepositions. It was not given the image and was instructed not to add new objects, actions, attributes, or scene information. Content-word lists were used because pilot regeneration from shuffled full descriptions often recovered text resembling the original narratives. This procedure was intended to restore sentence form while reducing reconstruction of the original organization. It does not ensure different image relations: the original image-derived words may still support plausible or image-consistent reconstructions. The examples also vary in fluency, so regeneration is an intended grammaticality control rather than a validated grammaticality-matched condition. We inspected random samples of the regenerated descriptions for additions beyond the supplied content words (Supplementary Table 1). The examples also show that grammaticality and fluency varied, and we do not report independent evaluations of image correspondence for the original or regenerated descriptions. Regenerated descriptions were embedded using the same text encoder and L2-normalization procedure as the other linguistic feature spaces.

Regenerated descriptions were produced from content-word lists rather than from full bag-of-words descriptions. In pilot tests, language models given shuffled full descriptions often reconstructed sentences resembling the original narratives, likely because the full word inventory, including function words, constrained the output. Using extracted content words was intended to reduce recovery of the original image-specific organization while retaining core scene-related vocabulary. It does not ensure that reconstructed relations differ from those in the image. This choice was also supported by the main encoding results, where bag-of-words and content-word controls did not differ significantly in their ability to predict brain responses. However, this does not establish equivalence between those inputs.

### Textual similarity between original and regenerated descriptions

To quantify textual similarity between original and regenerated descriptions, we compared all 73,000 aligned description pairs after lowercasing, punctuation normalization, and removal of function words. Lexical overlap was measured using multiset token F1, defined as the harmonic mean of token precision and recall. Local verbatim overlap was measured using exact bigram recall, exact trigram recall, and the longest exact contiguous shared sequence. Broader word-order preservation was measured using ROUGE-L, which is based on the longest common subsequence between two token sequences. These measures assess textual overlap rather than grammaticality or the correctness of depicted relations.

To test whether order-sensitive similarity exceeded what would be expected from the shared lexical inventory alone, we generated 1,000 within-pair null realizations by randomly reordering the regenerated non-function words within each pair. This null procedure preserved each description’s vocabulary, word count, repeated words, and original–regenerated pairing, while disrupting word order. Ninety-five percent confidence intervals were estimated by bootstrapping description pairs 5,000 times. One-sided Monte Carlo permutation probabilities were calculated using the plus-one correction and adjusted across the five order-sensitive metrics using the Holm procedure.

### Visual feature extraction

To compare self-supervised and language-aligned visual representations within a common broad architecture family and large-data regime, we used two Vision Transformer Large (ViT-L) models pretrained at internet scale: DINOv3 (Sim’eoni et al., 2025) and SigLIP2 (Tschannen et al., 2025). Both use very large web-derived training resources, but their image distributions and training recipes are not matched. Internet-scale training motivates the comparison without establishing that dataset differences are negligible.

- **Self-supervised visual model.** Visual features were extracted using DINOv3, which is trained without language supervision. We extracted a global image representation from the final transformer layer after standard preprocessing and applied L2 normalization.
- **Language-aligned vision model.** Language-aligned visual features were extracted using SigLIP2, trained with an image-text objective using a pairwise sigmoid loss. We extracted a global image embedding from the final transformer layer and applied L2-normalized prior to encoding analysis.

### Voxelwise encoding

To predict voxelwise BOLD responses from model-derived features, we fit ridge regression independently for each voxel and feature space, separately for each subject. For each feature space, we reduced dimensionality using principal component analysis (PCA), retaining 97% of variance. Stimuli were split into training (80%, N=8,000) and held-out test (20%, N=2,000) sets. Both the feature matrices and voxel responses were z-scored using the training set statistics.

The ridge regression regularization parameter values were selected via 7-fold cross-validation over 100 logarithmically spaced **α** values (10^-8^-10^10^). After selecting the optimal **α** per voxel, models were refit on the full training set and evaluated on the held-out test set using the coefficient of determination (R2).

### Variance partitioning

To quantify the unique contribution of a feature set A beyond a baseline feature set B, we implemented variance partitioning following standard approaches (e.g., (Wang et al., 2023)). A joint feature space (A+B) was constructed by concatenating raw features prior to PCA. We fit ridge models for the baseline (B) and joint (A+B) feature spaces to obtain (R^2^_B_), and computed unique explained variance as:

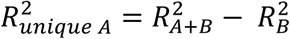

### Statistical tests

Statistical significance of both explained variance and unique variance was assessed via non-parametric bootstrap resampling of paired true and predicted test responses (2,000 iterations; resampling test images with replacement). Empirical p-values were defined as the proportion of bootstrap samples yielding variance estimates ≤ 0.

Voxelwise p-values were corrected for multiple comparisons using the Benjamini–Hochberg false discovery rate (FDR) procedure with q < 0.05.

### ROI-level analysis

For ROI summaries, we grouped NSD ROIs into five functional systems: early visual cortex (EVC), face-selective regions, place-selective regions, body-selective regions, and word-selective regions. Within each subject and ROI, we computed the mean unique variance across voxels. Group-level significance across subjects (N = 8) was assessed using two-sided Wilcoxon signed-rank tests against zero.

For the comparison of original- and regenerated-description gains beyond bag-of-words, each gain was defined as the held-out R² of the corresponding joint model minus that of the bag-of-words baseline. Subject-level regenerated gains were tested against zero using a two-sided sign-flip test across eight subjects. The difference between original and regenerated gains was tested using a two-sided paired subject-level sign-flip test. Effect estimates are reported with 95% confidence intervals. The regenerated-to-original gain ratio summarizes relative magnitude; it is not an estimate of shared explained variance. These comparisons were distinguished from the original-over-regenerated contrast, which used regenerated descriptions as the baseline.

### Image-wise prediction-difference alignment analysis

To characterize images associated with the model differences, we computed a prediction-difference alignment score for each test image using the observed response and the difference between joint-model and baseline-model predictions:

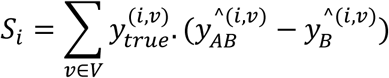

where

- 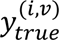 denotes the observed BOLD response
- *y_AB_* and *y_B_* are predictions from the joint and baseline models
- V denotes the set of voxels included in the analysis.

The term *y_AB_* − *y_B_* is the difference between joint-model and baseline-model prediction s. Thus *S_i_*summarizes alignment between this prediction difference and the observed response pattern. Because the models are separately fitted, the prediction difference should not be interpreted as a uniquely isolated feature component. The score is not a per-image R² or an exact decomposition of incremental R²: unlike a difference in squared prediction errors, it does not include the difference in squared prediction magnitudes.

Summation was restricted to voxels exhibiting significant unique variance (FDR correction, q < 0.05). For each model comparison, *S_i_* was computed for all 2,000 held-out test images and used to rank them. Positive and negative values indicate the sign of prediction-difference alignment, and near-zero values indicate little net alignment under this score; these labels do not directly describe changes in per-image prediction error. Because voxel inclusion was based on the same test-set analysis, the selected image groups are treated as exploratory.

### Image-level semantic annotation of stimulus groups

To characterize differences between high-score and near-zero images, we assigned three study-specific descriptive measures: compositional structure, semantic relations, and social interaction. Image groups were selected suing the only for post hoc characterization of the stimulus groups and were not included in the encoding models. The ratings are exploratory annotations rather than established benchmarks of compositionality. Independent inter-rater reliability is not reported.

The **compositional structure score** measured the extent to which multiple scene elements were organized into a coherent event or spatial layout. Scores ranged from 0 to 12 and were calculated as the sum of four ordinal subscores, each ranging from 0 to 3: agent multiplicity, action/interaction, spatial layout, and entity diversity. Agent multiplicity captured the number and organization of animate agents, ranging from no salient agent to multiple agents or a group. Action/interaction ranged from a static scene to a clear multi-agent or object-directed event. Spatial layout ranged from an isolated close-up to a scene with clear foreground–background organization and substantial spatial depth. Entity diversity ranged from a single object or entity type to multiple distinct agents, objects, and contextual elements. Higher scores therefore indicate richer organization of agents, actions, objects, and spatial context. Because agent multiplicity and entity diversity contribute to the composite, it describes scene richness as well as organization and does not isolate relations independently of entity content.

The **semantic relational score** measured the number of distinct, visually identifiable relations among salient entities. These included agent–action–object relations, agent–agent and human– animal interactions, object–object spatial or functional relations, and event–context relations. Repeated instances of the same relation were counted only when they represented distinct relational content. To prevent crowded scenes or repeated objects from disproportionately influencing the measure, scores were capped at 8. Higher scores indicate a greater number of distinct semantic relations.

The **social interaction score** measured the strength of socially meaningful interactions among animate agents on a scale from 0 to 3. A score of 0 indicated no visible social or agent–agent interaction; 1 indicated the presence of social agents with weak or ambiguous interaction; 2 indicated a clear dyadic or small-group interaction, such as conversation, competition, assistance, or directed human–animal interaction; and 3 indicated a rich, coordinated multi-agent event. Higher scores therefore indicate stronger and more explicit social interaction.

For each contrast and annotation measure, we compared the high-score and near-zero groups using two-sided Mann–Whitney U tests, with 20 images per group. In Fig. 3, individual image scores are shown as circles overlaid on box plots. Statistical significance is denoted by *p < 0.05, **p < 0.01, and ***p < 0.001.

## References

Aronson, S., Adkins, M. S., & Greene, M. R. (2025). Divergent roles of visual structure and conceptual meaning in scene detection and categorization. Journal of vision, 25(14), 21–21. 10.1167/jov.25.14.21

Bar, M. (2004). Visual objects in context. Nat Rev Neurosci, 5(8), 617–629. 10.1038/nrn1476

Bavaresco, A., Klarić, I., Fernández, R., & Moens, M.-F. (2026). What Makes Linguistic Representations Good Models of High-Level Visual Perception in the Human Brain? *arXiv preprint arXiv:2C07.1C214*.

Cavanagh, P. (2021). The Language of Vision. Perception, 50(3), 195–215. 10.1177/0301006621991491

Chen, T., Kornblith, S., Swersky, K., Norouzi, M., & Hinton, G. E. (2020). Big Self-Supervised Models are Strong Semi-Supervised Learners. *ArXiv*, *abs/200C.10029*.

Collins, J. A., & Olson, I. R. (2014). Knowledge is power: how conceptual knowledge transforms visual cognition. Psychon Bull Rev, 21(4), 843–860. 10.3758/s13423-013-0564-3

Conwell, C., Prince, J. S., Konkle, T., Isik, L., & Alvarez, G. A. (2024). Rethinking language-alignment in human visual cortex with syntax manipulation and word models.

Doerig, A., Kietzmann, T. C., Allen, E., Wu, Y., Naselaris, T., Kay, K., & Charest, I. (2025). High-level visual representations in the human brain are aligned with large language models. Nat Mach Intell, 7(8), 1220–1234. 10.1038/s42256-025-01072-0

Doerig, A., Kietzmann, T. C., Allen, E. J., Wu, Y., Naselaris, T., Kay, K. N., & Charest, I. (2024). Visual representations in the human brain are aligned with large language models. arXiv preprint arXiv:2209.11737,

Epstein, R. A., & Baker, C. I. (2019). Scene Perception in the Human Brain. Annu Rev Vis Sci, 5, 373–397. 10.1146/annurev-vision-091718-014809

Fairhall, S. L., & Caramazza, A. (2013). Brain regions that represent amodal conceptual knowledge. J Neurosci, 33(25), 10552–10558. 10.1523/jneurosci.0051-13.2013

Greene, M. R. (2013). Statistics of high-level scene context. Frontiers in Psychology, 4. 10.3389/fpsyg.2013.00777

Hafri, A., & Firestone, C. (2021). The Perception of Relations. Trends in cognitive sciences, 25(6), 475–492. 10.1016/j.tics.2021.01.006

Hafri, A., Trueswell, J. C., & Epstein, R. A. (2017). Neural Representations of Observed Actions Generalize across Static and Dynamic Visual Input. The Journal of Neuroscience, 37(11), 3056–3071. 10.1523/jneurosci.2496-16.2017

Isik, L., Koldewyn, K., Beeler, D., & Kanwisher, N. (2017). Perceiving social interactions in the posterior superior temporal sulcus. Proceedings of the National Academy of Sciences, 114(43), E9145–E9152. doi:10.1073/pnas.1714471114

Jackendoff, R. (1985). Semantics and Cognition. The Philosophical Review, 94(1), 111. 10.2307/2184719

Kominsky, J. F., & Wenig, K. (2025). Causal Perception(s). Cogn Sci, 49(9), e70107. 10.1111/cogs.70107

Lake, B. M., Ullman, T. D., Tenenbaum, J. B., & Gershman, S. J. (2017). Building machines that learn and think like people. Behavioral and Brain Sciences, 40, e253, Article e253. 10.1017/S0140525X16001837

Luo, A., Henderson, M., Wehbe, L., & Tarr, M. (2024). Leveraging Vision and Language Generative Models to Understand the Visual Cortex. Journal of vision, 24(10), 1333–1333. 10.1167/jov.24.10.1333

Malik, M., & Işık, L. (2023). Relational visual representations underlie human social interaction recognition. Nature Communications, 14(1). 10.1038/s41467-023-43156-8

Oliva, A., & Torralba, A. (2007). The role of context in object recognition. Trends Cogn Sci, 11(12), 520–527. 10.1016/j.tics.2007.09.009

Panchal, P., Polara, V., U, S., Baz, A., & Patel, S. K. (2026). Deep learning–driven image captioning: Progress through transformers and large language models. PLoS One, 21(3), e0345012. 10.1371/journal.pone.0345012

Peelen, M. V., Berlot, E., & de Lange, F. P. (2024). Predictive processing of scenes and objects. Nat Rev Psychol, 3, 13–26. 10.1038/s44159-023-00254-0

Pitcher, D., & Ungerleider, L. G. (2021). Evidence for a Third Visual Pathway Specialized for Social Perception. Trends Cogn Sci, 25(2), 100–110. 10.1016/j.tics.2020.11.006

Popham, S. F., Huth, A. G., Bilenko, N. Y., Deniz, F., Gao, J. S., Nunez-Elizalde, A. O., & Gallant, J. L. (2021). Visual and linguistic semantic representations are aligned at the border of human visual cortex. Nat Neurosci, 24(11), 1628–1636. 10.1038/s41593-021-00921-6

Radford, A., Kim, J. W., Hallacy, C., Ramesh, A., Goh, G., Agarwal, S., Sastry, G., Askell, A., Mishkin, P., Clark, J., Krueger, G., & Sutskever, I. (2021). Learning Transferable Visual Models From Natural Language Supervision. International Conference on Machine Learning,

Rajaei, K., Cichy, R. M., & Soltanian-Zadeh, H. (2026). VLMs using language-guided inference capture context-sensitivity of human object recognition behavior. Array, 29, 100731. 10.1016/j.array.2026.100731

Rong, B., Gifford, A. T., Düzel, E., & Cichy, R. M. (2025). The time course of visuo-semantic representations in the human brain is captured by combining vision and language models. In: eLife Sciences Publications, Ltd.

Sim’eoni, O., Vo, H. V., Seitzer, M., Baldassarre, F., Oquab, M., Jose, C., Khalidov, V., Szafraniec, M., Yi, S., Ramamonjisoa, M., Massa, F., Haziza, D., Wehrstedt, L., Wang, J., Darcet, T., Moutakanni, T., Sentana, L., Roberts, C., Vedaldi, A.,…Bojanowski, P. (2025). DINOv3.

Simanova, I., Hagoort, P., Oostenveld, R., & van Gerven, M. A. (2014). Modality-independent decoding of semantic information from the human brain. Cereb Cortex, 24(2), 426–434. 10.1093/cercor/bhs324

Tschannen, M., Gritsenko, A., Wang, X., Naeem, M. F., Alabdulmohsin, I. M., Parthasarathy, N., Evans, T., Beyer, L., Xia, Y., Mustafa, B., H’enaff, O., Harmsen, J., Steiner, A., & Zhai, X.-Q. (2025). SigLIP 2: Multilingual Vision-Language Encoders with Improved Semantic Understanding, Localization, and Dense Features. *ArXiv*, *abs/2502.1478C*.

Võ, M. L. (2021). The meaning and structure of scenes. Vision Res, 181, 10–20. 10.1016/j.visres.2020.11.003

Wang, A. Y., Kay, K., Naselaris, T., Tarr, M. J., & Wehbe, L. (2023). Better models of human high-level visual cortex emerge from natural language supervision with a large and diverse dataset. Nature Machine Intelligence, 5(12), 1415–1426.

Wischnewski, M., & Peelen, M. V. (2021). Causal neural mechanisms of context-based object recognition. Elife, 10, e69736.

Wurm, M. F., & Caramazza, A. (2019). Distinct roles of temporal and frontoparietal cortex in representing actions across vision and language. Nat Commun, 10(1), 289. 10.1038/s41467-018-08084-y

Ye, Q., Li, C., Zeng, X., Li, F., & Fan, H. (2025). Painting with Words: Elevating Detailed Image Captioning with Benchmark and Alignment Learning. 10.48550/arxiv.2503.07906

