## Supplementary Information for "Language-aligned model and structured scene descriptions reveal sensitivity to compositional scene in the high-level visual cortex"

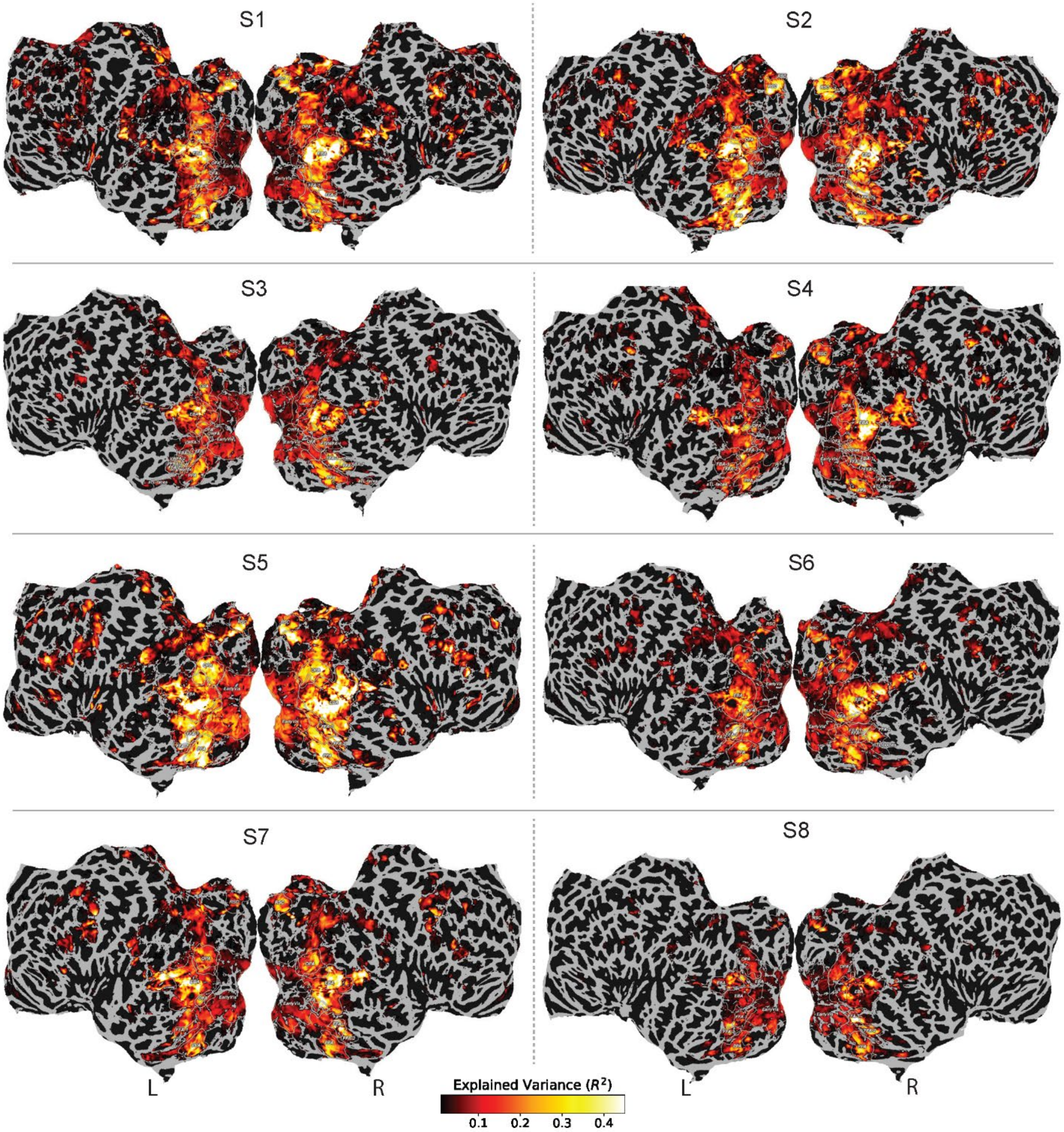

*Supplementary* ***Fig. 1.*** *Variance explained ($R^2$) by the full narrative description, shown on a cortical flatmap for all subjects. Only significant voxels are displayed (bootstrap test, $p < 0.05$, FDR-corrected). The text encoder is CLIP.*

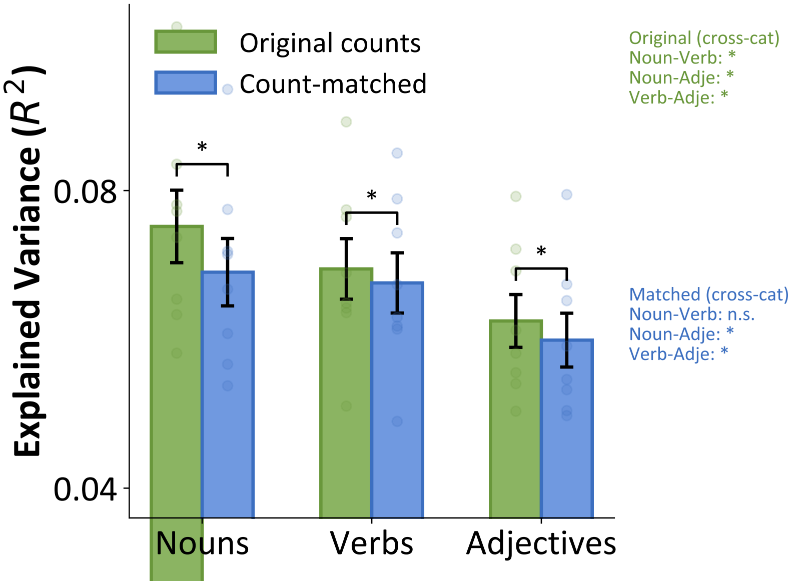


***Supplementary Fig. 2. Explained variance ($R^2$) across lexical categories in the original and count-matched data.*** *Green bars show results from the original word counts, and blue bars show results after matching word counts across categories to the minimum available count. Error bars indicate variability across comparisons (standard deviation), and points show individual values. Asterisks mark significant pairwise differences ($p < 0.05$, Wilcoxon signed-rank test, FDR-corrected). In the original-count condition, nouns showed the highest explained variance, followed by verbs and adjectives. After count matching, explained variance decreased overall; the difference between nouns and verbs was no longer significant, whereas adjectives remained significantly lower than both nouns and verbs.*

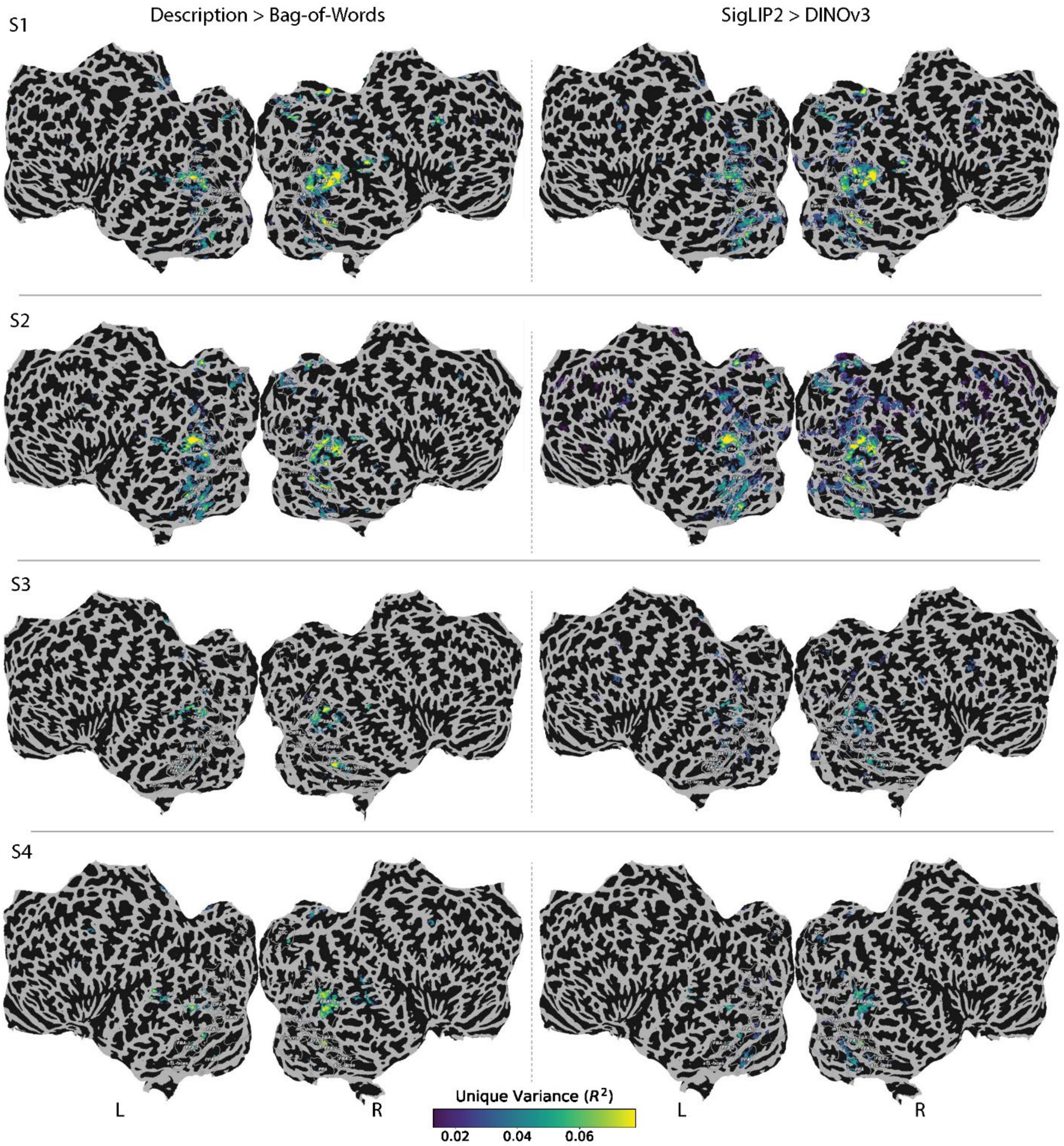
Description > Bag-of-Words
SigLIP2 > DINOv3
S1
S2
S3
S4
L
R
L
R
Unique Variance ($R^2$)
0.02
0.04
0.06

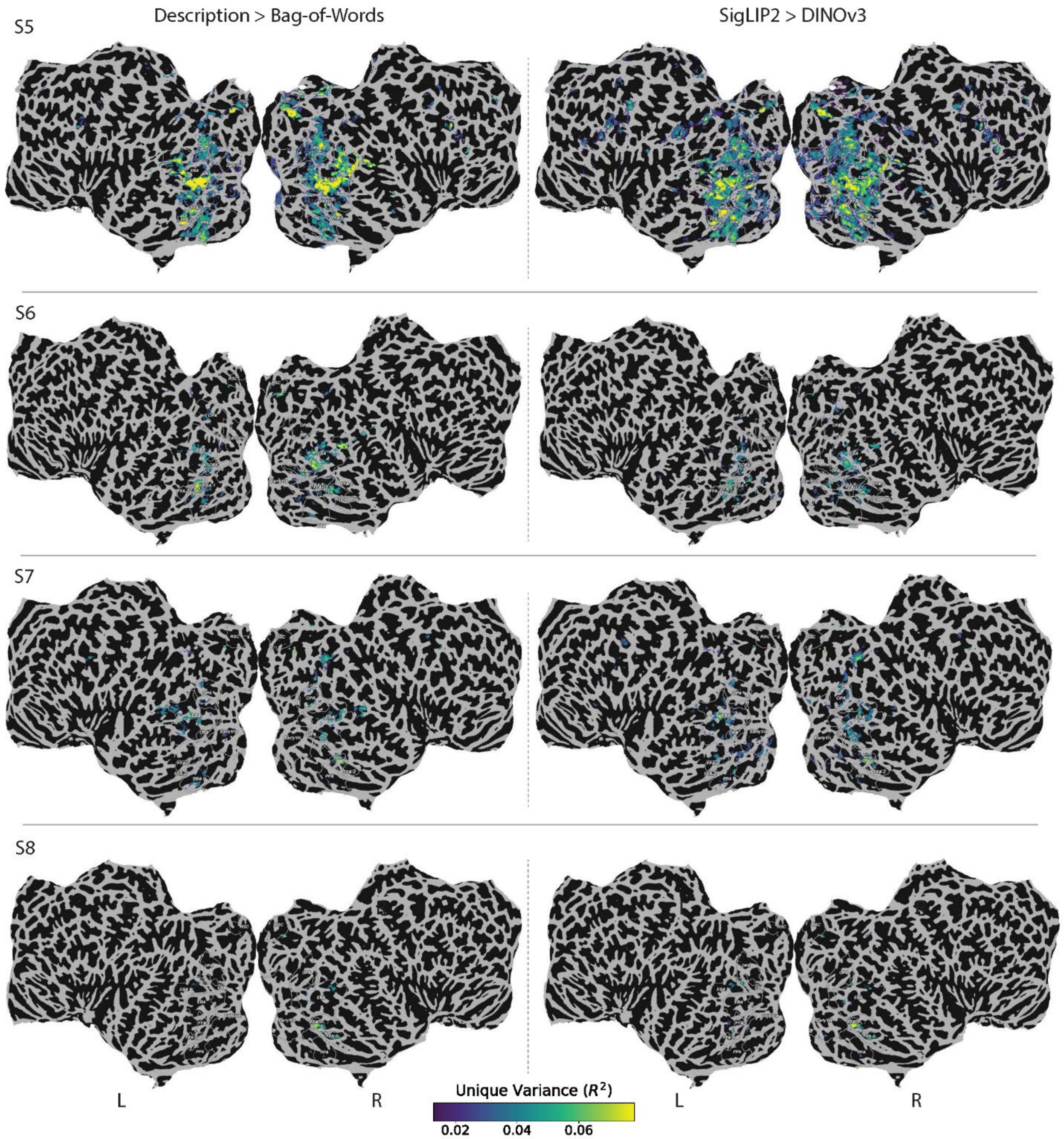


***Supplementary Fig. 3. All-subject flatmaps for the main unique-variance contrasts.*** *Voxel-wise unique explained variance ($R^2$) for narrative descriptions relative to bag-of-words (left) and for SigLIP2 relative to DINOv3 (right), shown for all subjects. In each joint encoding model, both feature sets were entered simultaneously; unique $R^2$ was defined as joint-model $R^2$ minus baseline-model $R^2$. Only significant voxels are displayed (bootstrap test, $p < 0.05$; FDR-corrected).*

***Supplementary Table 1. Examples of text controls used in the regenerated-description analysis.***

*Examples were randomly sampled using a fixed random seed, independently of encoding performance. Columns show original descriptions, extracted content-word lists, and regenerated descriptions. Regeneration used the content-word list without access to the image or original narrative, with instructions to use the supplied content-words and add only necessary function words.*

| Original Description | Content Words | Regenerated Description |
|---|---|---|
| A vibrant cityscape with modern skyscrapers reflecting sunlight, a riverboat glides on calm waters beneath a clear blue sky. | cityscape, waters, reflecting, blue, modern, glides, riverboat, skyscrapers, clear, sky, vibrant, calm, beneath, sunlight | Waters glides beneath blue modern clear and vibrant sky while reflecting with calm cityscape, riverboat, skyscrapers, and sunlight. |
| A vibrant lily blooms in a sleek black rectangular planter, surrounded by delicate green foliage against a softly lit window backdrop. | lit, black, green, delicate, rectangular, backdrop, blooms, surrounded, lily, softly, vibrant, window, planter, foliage, sleek | A planter is shown in lit black backdrop and green surrounded with delicate rectangular blooms, lily softly window, and vibrant sleek foliage. |
| A female tennis player in mid-swing on a court with an Emirates US Open Series backdrop. | female, court, tennis, player, emirates, backdrop, open, us, series, midswing | A tennis court is midswing in open, emirates, backdrop, and us with female player and series. |
| Two oxen harnessed to a cart on a rural road under a clear sky with people in the background. | oxen, two, background, clear, road, harnessed, sky, rural, cart, people | Two oxen are harnessed under clear, background, road, and sky with rural cart and people. |
| Stacked luggage on metal shelves in a store with colorful patterns and black suitcases. | stacked, colorful, black, suitcases, store, shelves, metal, luggage, patterns | Colorful black metal suitcases are stacked in store and patterns with shelves and luggage. |

| | | |
|---|---|---|
| A bluebird perches on a bare branch against a blurred, overcast sky. | sky, bare, blurred, perches, overcast, branch, bluebird | A bare overcast bluebird perches under the sky while blurred with branch. |
| A couple poses for a photo in formal attire against an ornate wall backdrop. | formal, couple, wall, backdrop, poses, attire, photo, ornate | A couple poses wearing formal attire in wall and backdrop with ornate photo. |
| A close-up of a pizza with melted cheese and mushrooms on a dark wooden surface, illuminated by a warm candle in the background. | cheese, illuminated, surface, melted, background, pizza, dark, candle, warm, mushrooms, wooden, closeup | A melted dark warm surface is illuminated in wooden background with cheese, pizza, candle, mushrooms, and closeup. |
| A female tennis player in action on clay court, preparing to hit a forehand with a pink backdrop behind her. | player, court, her, backdrop, hit, female, behind, pink, forehand, clay, tennis, preparing, action | A tennis player hit behind pink court and backdrop while preparing with female forehand her and clay action. |
| A vintage-style clock with neon 'Melrose' sign on a weathered building facade. | Vintage, style, clock, neon, weathered, facade, building, melrose, sign | A clock is weathered in building and sign with vinta gestyle, neon, facade, and melrose. |

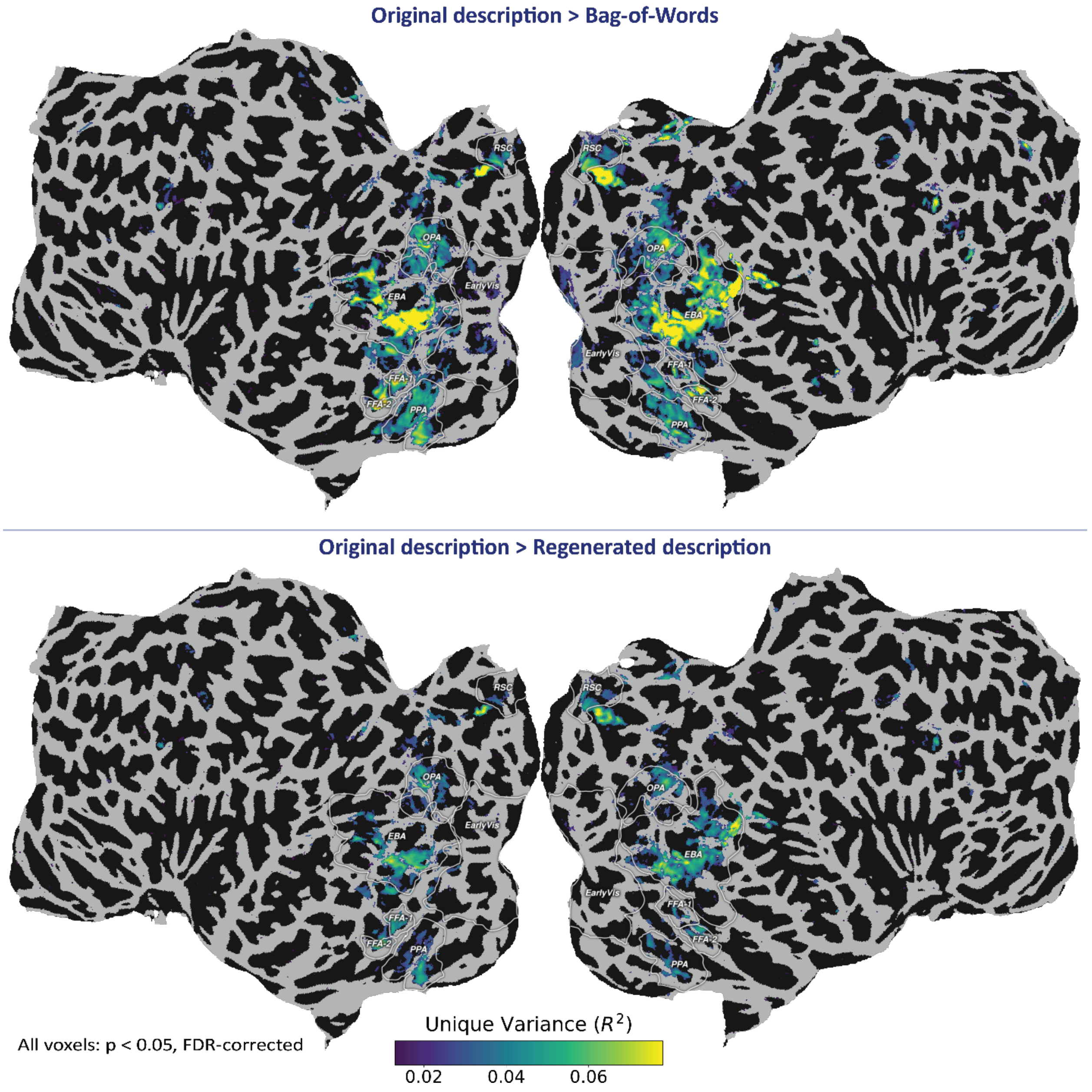


***Supplementary Fig. 4. Cortical maps for the regenerated-description control in subject S5.*** *Flatmaps show voxel-wise unique explained variance for original narrative descriptions relative to bag-of-words and regenerated descriptions. Regenerated descriptions were produced from the same content-word lists as the original descriptions, with only necessary function words added. Both maps use the same color scale and show only significant voxels. The regenerated-description contrast was weaker than the bag-of-words contrast but had a similar concentration in high-level visual cortex (Fig. 2c).*

***Supplementary Table 2. Textual similarity between original and regenerated descriptions across 73000 aligned description pairs.***

| Measure | Observed result | Null/comparison | Interpretation |
|---|---|---|---|
| Non-function-token overlap F1 | 0.957 [0.956, 0.957] | — | High lexical overlap |
| Longest exact shared sequence | Median = 2 tokens | 11.3% ≥3; 1.6% ≥4 | Long verbatim copying was uncommon |
| ROUGE-L | 0.446 [0.445, 0.446] | Null = 0.405 [0.404, 0.406] | Small retained word ordering |
| ROUGE-L difference | 0.041 [0.040, 0.041] | ($p_{Holm}$=0.01) | Greater than expected from shared vocabulary |

***Supplementary Table 3. Subject-wise Pearson correlations between voxel-wise unique $R^2$ values for Description > Bag-of-Words and SigLIP2 > DINOv3.*** *Statistics are reported for all analyzed voxels and separately for each ROI. Values are reported as sample size (n), Pearson correlation coefficient (r), and corresponding p-value.*

| ***Subject*** | ***All Voxels (n, r, p)*** | ***Bodies*** | ***Faces*** | ***Places*** | ***EVC*** | ***Other*** |
|---|---|---|---|---|---|---|
| ***S01*** | 29,489; **0.58**; <0.001 | 8,851; **0.74**; <0.001 | 679; **0.74**; <0.001 | 4,333; **0.33**; <0.001 | 3,196; **0.12**; <0.001 | 12,430; **0.41**; <0.001 |
| ***S02*** | 48,969; **0.74**; <0.001 | 10,909; **0.78**; <0.001 | 1,582; **0.77**; <0.001 | 7,173; **0.58**; <0.001 | 3,297; **−0.12**; <0.001 | 26,008; **0.60**; <0.001 |
| ***S03*** | 12,201; **0.53**; <0.001 | 6,072; **0.49**; <0.001 | 498; **0.42**; <0.001 | 1,049; **0.02**; 0.594 | 413; **−0.08**; 0.125 | 4,169; **0.48**; <0.001 |
| ***S04*** | 9,054; **0.44**; <0.001 | 2,960; **0.62**; <0.001 | 851; **0.74**; <0.001 | 1,502; **0.12**; <0.001 | 390; **0.00**; 0.988 | 3,351; **0.28**; <0.001 |
| ***S05*** | 59,555; **0.71**; <0.001 | 16,019; **0.73**; <0.001 | 2,349; **0.67**; <0.001 | 8,497; **0.64**; <0.001 | 6,715; **−0.12**; <0.001 | 25,975; **0.69**; <0.001 |
| ***S06*** | 11,502; **0.44**; <0.001 | 5,841; **0.52**; <0.001 | 848; **0.68**; <0.001 | 740; **0.02**; 0.523 | 539; **−0.34**; <0.001 | 3,534; **0.14**; <0.001 |
| ***S07*** | 10,447; **0.40**; <0.001 | 3,908; **0.40**; <0.001 | 152; **0.82**; <0.001 | 1,020; **0.14**; <0.001 | 1,005; **−0.55**; <0.001 | 4,362; **0.42**; <0.001 |
| ***S08*** | 2,095; **0.62**; <0.001 | 1,043; **0.36**; <0.001 | 417; **0.88**; <0.001 | 37; **0.34**; 0.042 | 104; **−0.02**; 0.841 | 494; **0.55**; <0.001 |

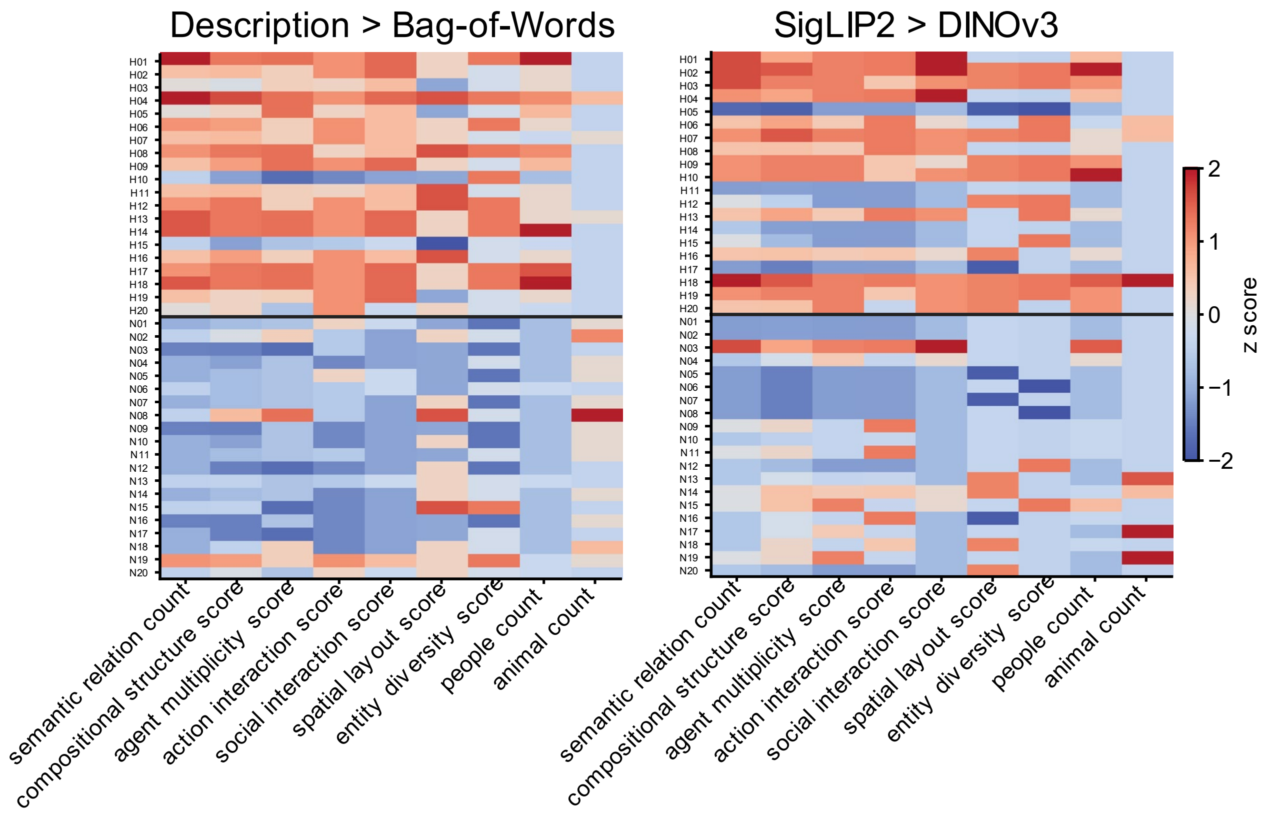


***Supplementary Fig. 5. Exploratory per-image semantic ratings for the stimulus-level analyses in Fig. 3.*** *Heatmaps show author assigned semantic annotations for the high-score and near-zero image sets in the Description > Bag-of-Words and SigLIP2 > DINOv3 contrasts. Rows correspond to individual images, ordered as in Fig. 3, and columns correspond to semantic metrics, including semantic relational score, compositional structure score, agent multiplicity, action/interaction score, social interaction score, spatial layout score, entity diversity, people count, and animal count. Values are z-scored separately within each measure for visualization, so larger values indicate higher ratings relative to the other images in the same analysis. The horizontal separation marks the boundary between high-score and near-zero groups. The heatmaps describe group difference and variability under this annotation scheme; they do not establish relational selectivity independently of entity content or scene richness.*
